# Weakest link epistasis and the geometry of genetic load

**DOI:** 10.64898/2025.12.08.693057

**Authors:** Florian J.F. Labourel, Florence Bansept, David M. McCandlish

## Abstract

Because natural selection must optimise multiple traits at once, previous work has suggested that phenotypic dimensionality can substantially worsen the equilibrium fitness defect of a population relative to the phenotypic optimum. However, it remains unclear how conclusions drawn from classical theoretical phenotype–fitness maps extend to models grounded in explicit biological mechanisms. Here we introduce weakest-link epistasis (WLE), a framework in which fitness is determined by the least-fit phenotypic component, an extreme form of diminishing returns epistasis. We show that in this framework, increasing dimensionality amplifies the load in a manner comparable to, but surprisingly not more than, Fisher’s geometric model (FGM). Building on this similarity, we demonstrate why genetic load is often invariant across different rules for combining trait-specific fitness components into an overall organismal fitness. We explore these ideas by considering the family of models where the organismal fitness is determined based on the *ℓ*^*p*^-norm of the vector of trait-specific fitness defects, a framework that includes both FGM and WLE, but also captures a continuum of genetic architectures, ranging from generalist to specialist regimes. Altogether, our approach proposes a new perspective on the geometry of adaptive landscapes, and may help provide quantitative insight into the cost of complexity.

## 1 Introduction

The difference between the stationary mean fitness of a population and the fittest possible genotype is called the genetic load [1–4]. Since alleles are effectively neutral when their selection coefficient *s* << 1/*N*_*e*_, where *N*_*e*_ denotes the long-term (effective) population size of a species, the genetic load in a single-trait model is classically predicted to scale as 1/*N*_*e*_ [4–7]. However, this apparent predictability of the genetic load is complicated by the genetic architecture underlying fitness (e.g. supply of deleterious mutations [3], degree of epistasis [8]). Notably, increasing the number of fitness-contributing traits can dramatically amplify the supply of deleterious mutations and hence the genetic load [2, 9–11], thereby creating a mutational burden that scales with the complexity of organisms [12–14].

This burden of complexity has most often been studied through the lens of Fisher’s geometric model (FGM), under which the fitness landscape contains a single optimum in the phenotypic space. In this model, the genotypic load carried by an organism increases with the distance to the optimum of its phenotype mapped into the fitness space [10, 14, 15]. In FGM, complexity is thus traditionally captured by the dimensionality of the fitness landscape, with the assumption that each dimension could potentially represent a component (eg., a gene, a trait) contributing to fitness [14]. In this scenario, the genetic load was shown to be substantially exacerbated (relative to the single-trait case), to the extent that it can become proportional to the number of fitness-contributing components [2, 9].

Yet, if FGM has received some theoretical support [14, 16], it remains unknown how accurate and broad a description it is of *in vivo* fitness landscapes [17–20]. Indeed, FGM relies on a very specific assumption about how phenotypic traits combine to produce fitness—namely, that fitness depends on the Euclidean distance to the peak of their respective deviation from their optimum value— which calls into question the universality of its predictions, including those about the genetic load. Accordingly, FGM was extended to demonstrate the genetic load invariance across a much broader range of theoretical fitness models [12]. A central limitation of this finding, however, is that neither fitness nor the underlying definition of complexity is grounded with a clear biological rationale, raising questions about how complexity should be measured [12, 13, 21].

By contrast, a relevant yet unexplored principle governing the combined effect of traits on fitness is Liebig’s law of the minimum [22–24]. According to this law, an organism’s growth is simply dictated by the most limiting nutrient in its environment, that is its “weakest link”, much like the capacity of a barrel composed of staves of unequal length would be limited by its shortest stave. Though originally proposed (and criticised) for the response of organisms to resources [22], this concept conceals a significant potential for generalisation, especially at the cellular scale [24]. In particular, because biosynthetic molecules can either be taken up in the environment or produced through *de novo* synthesis, it is natural to extend this weakest link logic to the mapping of metabolic traits to fitness such that fitness would be dictated by the least-fit trait [25]. Imagine a two-enzyme pathway, for instance, where one enzyme would be inefficient. In this scenario, the other enzyme would have little influence on the metabolic flux (and hence on fitness, provided fitness is positively correlated with the flux), because it has virtually no control on the flux. This epistasis-driven diminishing returns is the classical expectation from metabolic control theory [26, 27], where the flux increase due to enhancing an enzyme’s efficiency gets inescapably limited by the presence of other enzymes in the pathway [19, 28–31]. Notably, a similar mapping of traits onto fitness recently helped explain why some metabolic features may be far off their optimum [32]. It is also consistent with the existence of lethal mutations that make fitness equal zero regardless of how fit most of an organism’s genes are [33].

Based on this “law of the minimum”, we first describe the population genetics consequences of weakest link epistasis (WLE) in terms of fitness landscapes and mutant selection coefficients [25]. We then determine the genetic load under WLE and show that phenotypic dimensionality *n* amplifies the bias toward maladaptive traits *b*, which reflects the tendency of mutations to produce deleterious trait changes, yielding an effective maladaptive trait bias *nb*. To demonstrate the generality of this finding, we prove that a broad class of models for combining trait-specific fitness defects into an overall organismal fitness defect lead to the same genetic load, which we show is a consequence of the geometric relationships between equal fitness contours. We explore these results by considering a class of models where the organismal fitness defect is equal to the *ℓ*^*p*^ norm of the vector of trait-specific fitness defects, and discuss the extent to which these models can capture different phenotypic strategies (e.g. generalists versus specialists) despite the genetic load being identical.

## 2 Model & Results

Consider a haploid population of individuals that have a number *n* of traits (or phenotypic dimensions). For a genotype *g*, let **F**_**g**_ = (*F*_*g*,1_, …, *F*_*g,d*_, …, *F*_*g,n*_), be the vector of corresponding fitness components where *F*_*g,d*_ denotes the trait-dependent fitness component associated with the value for the *d*-th phenotypic trait of genotype *g*. Each such fitness component captures the contribution of a trait to fitness (e.g. the fitness contribution brought by a metabolic reaction, by the shape of an organism, etc.). These fitness components are then combined through a function *f*_*org*_(**F**_**g**_) to yield the fitness *f*_*g*_ of an organism with genotype *g*:

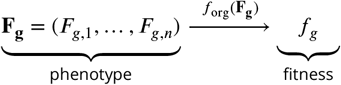

We consider the fitness components *F*_*g,d*_ to range from 0 to 1 so that values are easily comparable across traits and can be interpreted as a mapping of trait values into a bounded fitness space. Specifically, we define *F*_*g,d*_ = *f*_*org*_(1, 1, …, *F*_*g,d*_, …, 1) when all but component *d* of genotype *g* has been set equal to 1, i.e. the fitness component *F*_*g,d*_ has the interpretation that it is the fitness that would be conferred by the *d*-th trait of genotype *g* if all its other traits were set to their optimal values.

Based on these premises, we now provide a mathematical formalisation of weakest link epistasis and its direct population genetics consequences.

### 2.1 Weakest link epistasis from a population genetics perspective

In order to formalize the weakest link logic, we can define the fitness of a genotype *f*_*g*_ = *f*_*org*_(**F**_**g**_) as the minimum of the trait-dependent fitness components (see Figure 1):

**Figure 1.**
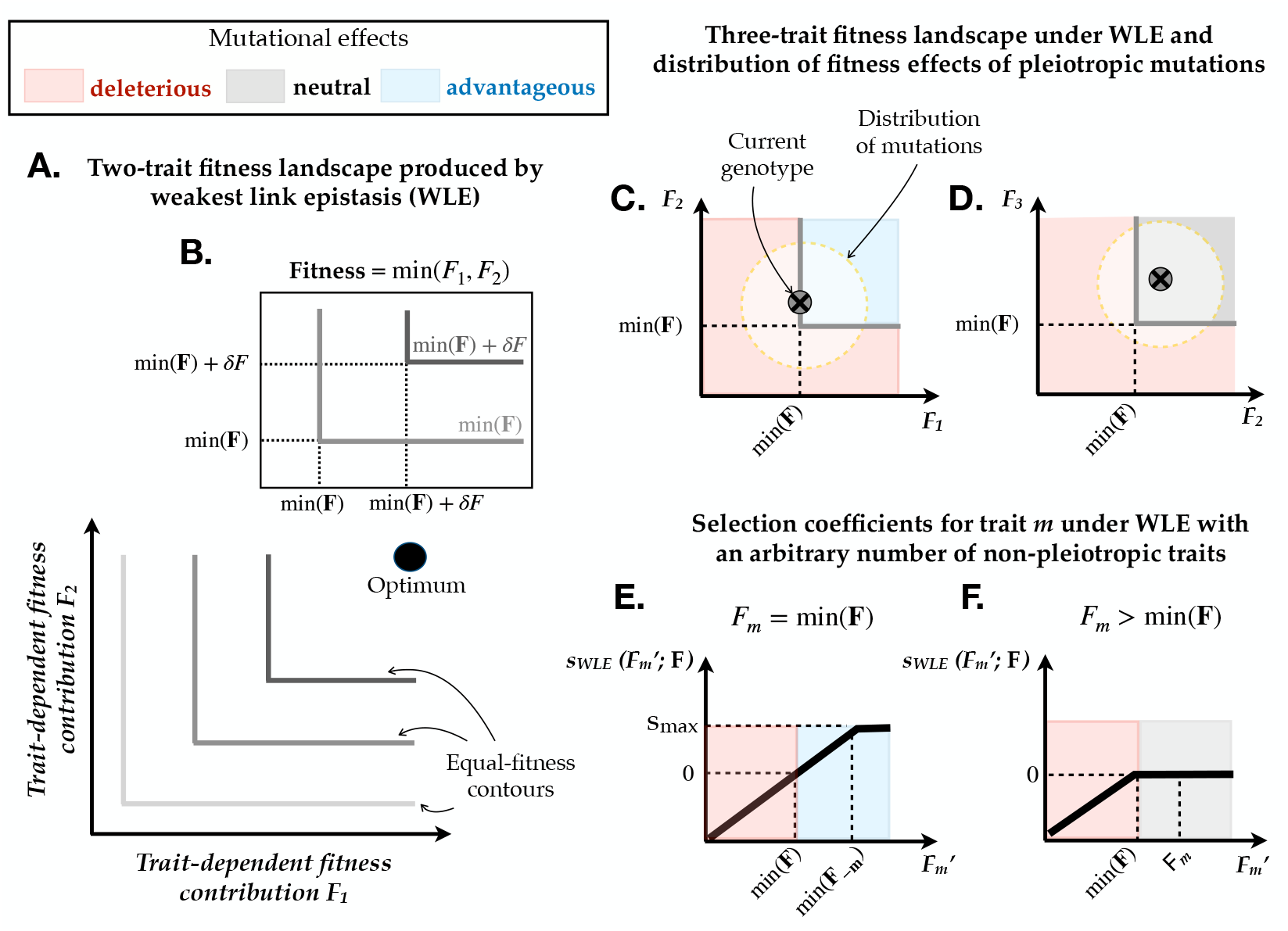
Explanation of the mechanism of weakest-link epistasis (WLE) and its population-genetic consequences when each trait *d* produces a fitness component *F*_*d*_. **A**–**B**. Panels A and B illustrate how WLE operates in a two-dimensional trait space where fitness is bounded (e.g, between 0 and 1) and reaches its optimum at the upper-right corner (black circle). In **B**, the contours of equal fitness are represented in shades of grey, with their values indicated above each contour. Under WLE, fitness equals the minimum of the fitness components, min(**X**): it thus remains equal to *F*_1_ as long as *F*_2_ > *F*_1_, and symmetrically for *F*_1_. Fitness increases by *δF* only when both fitness components increase by at least *δF* . In **A**., the resulting fitness landscape is composed of quarter-square contours centered on the optimum (in grey). **C**–**D**. Panels C and D illustrate why WLE produces an excess of neutral and deleterious mutations for the WLE fitness model when pleiotropic mutations affect two traits at a time. As an illustration, the mutational neighbourhood is delimited by circles (with orange-dotted boundaries). In **C**, when one of the mutated trait is the resident’s weakest link (trait 1 in this example), the resident genotype (grey cross-in-circle) lies on the equal-fitness contour (grey line). A range of mutations increasing this least-fit trait are advantageous (blue region, upper-right), yet comparatively more mutations are deleterious (red region). The exact balance of advantageous vs. deleterious mutations depends on the exact position of the other trait. In **D**, when mutations affect two traits that are not the resident’s weakest link, their effect is either neutral because the weakest link remains unchanged (grey region above the fitness contour) or deleterious otherwise (red region); no mutation can be advantageous in this case. **E**–**F**. Selective coefficients of non-pleiotropic mutations affecting only trait *m*. In **E**, for a mutation affecting the weakest link, the selection coefficient *s*_WLE_ increases (up to *s*_*max*_) until 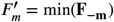, above which trait *m* is no longer the weakest link. In **F**, as in panel D, no mutation can be advantageous since the mutated trait *m* is not the weakest link.

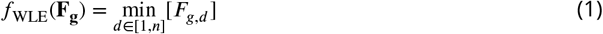

Because *F*_*g,d*_ only prescribes fitness when it is the minimum, or limiting factor, it represents the maximum fitness possible given that trait value (e.g., the maximum growth rate that ATP production or nucleotides synthesis can sustain).

#### 2.1.1 Selection coefficients under weakest link epistasis

If mutations are rare enough, a new mutant with genotype *g*^′^ is either fixed or lost before the next new mutation arises, which implies the absence of clonal interference [5, 7, 34]. We can thus drop genotype indexes and focus on a mutant with trait-dependent fitness components **F**^′^ and a resident with trait-dependent fitness components **F**. Hence, the selection coefficient (such that *f*_*g*_′ = *f*_*g*_(1 + *s*)) reads:

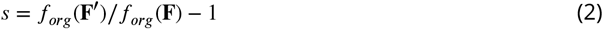

Under WLE, this implies that the selection coefficient of a mutant **F**^′^ is thus:

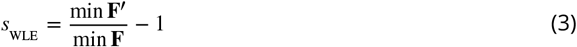

Whether mutations are deleterious (*s*_WLE_ < 0), neutral (*s*_WLE_ = 0) or advantageous (*s*_WLE_ > 0) there-fore simply depends on the sign of the difference between the weakest link of the mutant min (**F**^′^) and the weakest link of the resident min (**F**).

Yet, because many mutations are masked by the weakest link effect on fitness, selection is more complex and counterintuitive than it would be in the absence of epistasis. Crucially, this fitness model tends to inflate the supply of purely neutral mutations, since any mutation that preserves min (**F**^′^) = min (**F**) leads to *s*_WLE_ = 0. This happens whenever the weakest link component remains intact after mutation. The resulting extended range of neutral mutations includes any mutation that does not decrease any of the mutant’s fitness contributions below the weakest link value of the resident.

To further understand the selective effect of mutations, we can focus on the non-pleiotropic scenario where mutations only affect one trait at a time. Any mutant genotype *g*^′^ thus differs from its resident progenitor *g* by only one phenotypic trait value. We denote this mutant trait value 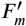 (in the fitness components space) when it affects trait *m*, so that 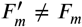. As mutants and residents share the same genotypic (and phenotypic) background **F**_**−m**_, the trait-dependent fitness components are all equal but for the mutant trait:

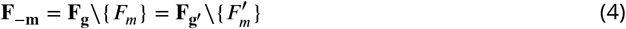

Weakest link epistasis creates a selective asymmetry between traits that becomes striking in the absence of pleiotropy. In particular, advantageous mutations are fully restricted to cases in which the mutated trait *m* is the resident’s weakest link, that is *F*_*m*_ = min(**X**). Provided that 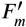 remains the mutant’s weakest link 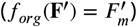, the selection coefficient reduces to:

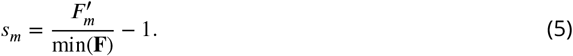

Otherwise, the (unchanged) second-worst trait becomes the mutant’s new weakest link. The mutant’s selective advantage thereby reaches an upper bound dictated by the second worst trait min(**F**_**−m**_), above which 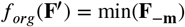, despite 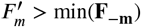. In such cases:

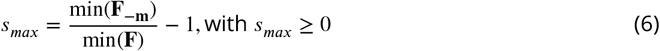

By contrast, many mutations reducing fitness components are latent and thus invisible to selection (*s*_WLE_ = 0); this can happen for any trait except the resident’s weakest link, provided that the decrease is small enough to leave the weakest link unchanged—that is, 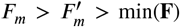 Otherwise, if the mutated trait becomes the new weakest link with 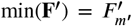 the mutation is deleterious, with a selection coefficient given by Eq. (5).

Altogether, the adaptive potential of 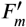 can be captured through a selection coefficient 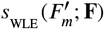 that only depends on itself and the resident set of trait-dependent fitness components **F**. Based on Eq. (3), this can be formally written as:

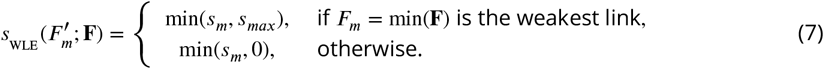

This expression captures both the mutational asymmetry between traits and the intrinsic limitation on advantageous mutations.

As pleiotropy increases, the mutational logic becomes progressively shifted away from pure neutrality. We can illustrate this phenomenon with a simple combinatorial scenario in which mutations affect subsets of traits. In particular, consider a mutation that affects *k* out of *n* traits of an organism (*k* ≤ *n*), where the subset of mutated traits is drawn at random from all 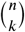 subsets (i.e. no modular pleiotropy). The degree of pleiotropy can thus be characterised as the proportion *k*/*n* of traits affected by a mutation. If each trait experiences the same mutation rate, the fraction of mutations affecting the weakest-link is simply given by 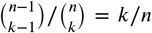, since mutations affecting the weakest-link trait can be combined with any *k* − 1 of the remaining *n* − 1 traits. With three traits, for example, two-dimensional mutations can either involve the weakest link together with one of the other traits (which occurs twice), or involve the two fittest traits (which occurs only once).

Because mutations evolving under pure neutrality rely on the weakest link remaining intact, the fraction of such mutations decreases proportionately with increasing pleiotropy, according to 1 − *k*/*n*. Under more universally pleiotropic scenarios, what instead becomes inflated is the supply of deleterious mutations: indeed, any pleiotropic mutation that brings at least one trait below the resident’s weakest-link value will be deleterious (the occurrence of cases D. in Figure 1 decreases in favour of cases C.).

Overall, the weakest-link effect (WLE) imposes a sharp limitation on the supply of advantageous mutations, and we now turn to its implications for the genetic load.

#### 2.1.2 Genetic load under weakest link epistasis

By removing unfit genotypes, natural selection biases the observed distribution of fitness, theoretically driving it to the optimum. However, mutations recurrently introduce deleterious genotypes that can segregate and even fix through random drift—despite being disfavoured—thereby limiting fitness improvement. This mutation–selection–drift balance leads to an evolutionarily stationary state for the distribution of fitness *ρ*^∗^(*f* ) whose mean deviates from the optimum. The resulting extent to which the evolutionary process is suboptimal can be quantified through the stationary genetic load *L*^∗^, which corresponds to the relative difference 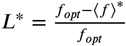 between mean stationary fitness ⟨*f*⟩^∗^ and the fitness optimum *f*_*opt*_.

One key determinant of the genetic load is the distribution of fitness over all possible genotypes (in the absence of selection), whose density we write as *ρ*(*f* ). To investigate the impact of WLE on the genetic load, we thus need to evaluate first how it influences *ρ*(*f* ). For that purpose, we assume trait-dependent fitness components to be independent and identically distributed random variables so that the genotypic density *ρ*(*F*_*d*_ ) is known for each *F*_*d*_ . The genotypic density of fitness *ρ*(*f* ) can then directly be obtained by combining the respective densities *ρ*(*F*_*d*_ ) with the fitness model *f*_*org*_(**F**). Under WLE, *f*_*org*_(**F**) = min(**F**) such that *ρ*_WLE_ (*f* ) is given by the distribution of the minimum of the set **F**. For analytical tractability, we thus choose a distribution for fitness components whose first order statistic *F*_(1)_ = min {*F*_1_, *F*_2_, …, *F*_*n*_} is tractable. The Beta distribution, when applied uniformly across traits such that each *F*_*d*_ obeys *ρ*(*F*_*d*_ ) ∼ Beta(*a, b*), matches this requirement when its first parameter is set to *a* = 1, which we discuss in more depth in Appendix B.1.1 in relation to the Kumaraswamy distribution (see also [35, 36]). Hence, the density of fitness components reads:

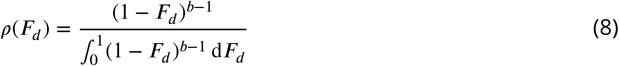

Here, the parameter *b* creates a bias toward maladaptive phenotypes on each trait by increasing the number of genotypes leading to low-fitness values. From now on, we refer to this as the maladaptive bias. Since 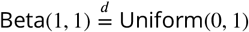, the unbiased case occurs when *a* = *b* = 1, and we can capture the strength of this per-trait maladaptive bias using log(*b*) (see Figure 2-A, assuming *n* = 1). Because we regard fitness as being the minimum of the *n* fitness components, the distribution of the first order statistics *ρ*(*F*_(1)_) for a sample of *n* independent draws from this distribution directly gives the genotypic fitness density under WLE (see Appendix B.1.1 for a proof):

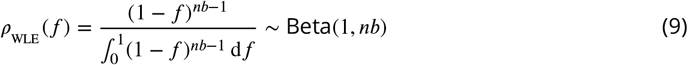

**Figure 2.**
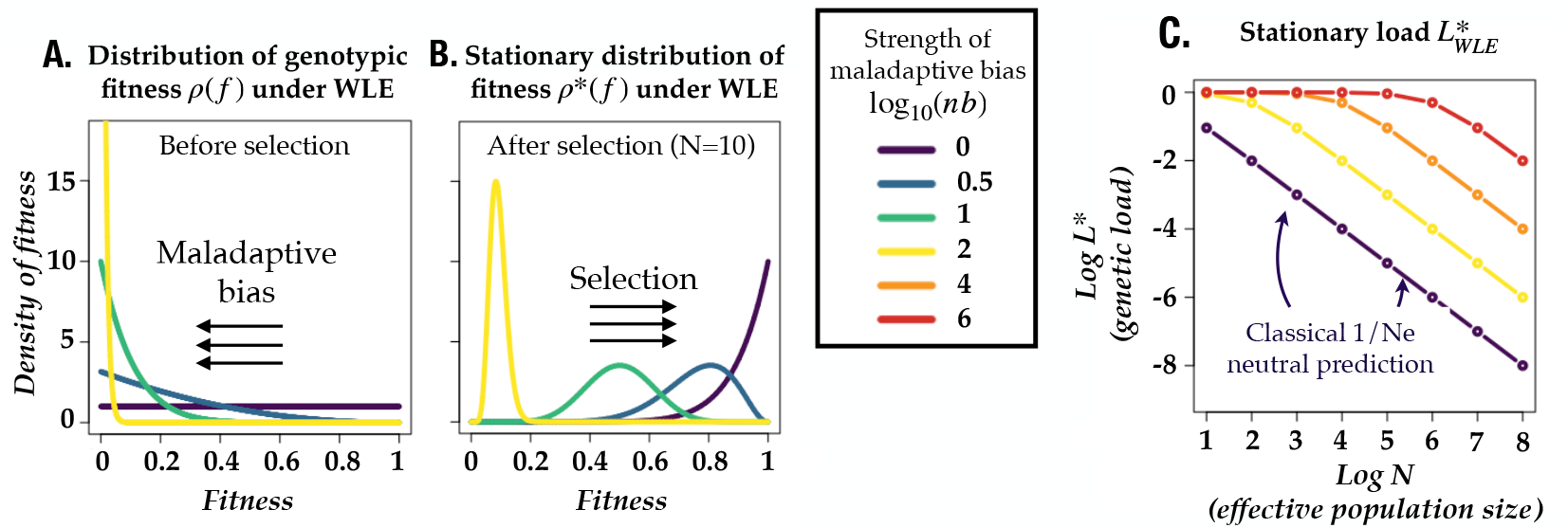
Distribution of fitness and evolution of maladaptation under weakest-link epistasis. Results are shown for varying levels of effective maladaptive biases, ranging from no bias (*log*(*nb*) = 0, dark blue) to strong bias (*log*(*nb*) = 6, red). **A**. Distribution of genotypic fitness *ρ*_WLE_ (*f* ) expected in the absence of selection. When each of the *n* trait-dependent fitness components follows Beta(1, *b*), genotypic fitness is distributed as Beta(1, *nb*). In this scenario, the per-trait maladaptive bias *b* and the number of traits *n* combine symmetrically to push the distribution toward lower fitness values. **B**. Evolutionarily stationary distribution of fitness 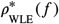 after selection (illustrated for N=10). Fitness still obeys a Beta distribution, Beta(*N, nb*), but compared to panel A, the distribution is shifted toward higher fitness by selection. **C**. Evolutionarily stationary load *L*^∗^ for different effective population sizes *N*, plotted on log-log scales. The classical population genetics prediction for the genetic load of a single trait scales as 1/*N* and appears as a straight decreasing line with slope −1. Increasing the effective maladaptive bias, either through a larger number of traits *n* or through a stronger intrinsic bias *b*, causes substantial maladaptation (*logL*^∗^ ≈ 0 so that *L*^∗^ → 1) even for large population sizes.

Remarkably, this expression highlights how dimensionality *n* mirrors the role of the maladaptive bias *b*, which motivates defining *nb* as an effective maladaptive bias (see Figure 2-A).

To gain insight into the genetic load, we can focus on the stationary state reached under the weak mutation regime, where a mutant genotype either gets fixed or lost [4, 12, 37, 38]. In this scenario, the evolutionary dynamics can be captured by a Markov chain where the transition rates from genotypic state *g* to *g*^′^ is simply given by the product of the mutation rate *μ*_*gg*_′ and the probability of fixation *P*_fix_(*g*^′^; *g*) of *g*^′^ in a population where *g* is the resident. Since this Markov chain can be shown to be reversible, it obeys detailed balance at stationarity, meaning forward and backward substitutions between two mutationally connected genotypes (from *g* to *g*^′^, and in reverse) occur at the same rate [4, 37]. Extending this detailed balance principle to all mutational neighbours (accessible through 1 mutational step), there exists a tractable solution for the stationary density *ρ*^∗^(*f* ) of the Markov chain owing to the link between *P*_fix_(*g*^′^; *g*), and the resident and mutant fitness values *f*_*g*_ and *f*_*g*_′ . Under the haploid Moran scenario and pairwise symmetrical mutations *μ*_*gg*_′ = *μ*_*g*_′_*g*_, this stationary fitness distribution has density [4]:

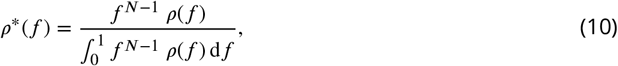

where *N* denotes the effective population size, which controls how strongly selection favours higher-fitness genotypes.

Using the density function of beta distributions, we can now plug Eq. (9) into Eq. (10) to obtain the stationary distribution for fitness under WLE. The resulting stationary distribution of *f* is distributed as Beta(*N, nb*):

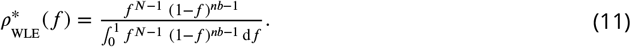

This expression also clearly shows the intuitive antagonistic effect between *N* and the effective bias *nb* (see Figure 2-B). This can be further formalised by deriving mean stationary fitness ⟨*f*⟩^∗^ as the first moment of the stationary distribution, which also coincides with the ratio between the *N*th and (*N* − 1)th moments of the distribution of genotypic fitness implied by Eq. (9). In our fitness space with upper bound 1, the stationary genetic load reduces to *L*^∗^ = 1 − ⟨*f*⟩^∗^, and hence WLE leads to:

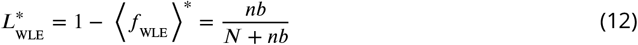

The saturating function in Eq. (12) shows that the genetic load scales linearly with *nb*/*N* when *N* ≫ *nb*, while otherwise the expression indicates significant maladaptation, especially in the hypothetical case where *nb* would be greater than *N* (see Figure 2-C). Note that, when *nb* and *N* are comparable, the stationary fitness distribution is akin to a Gaussian distribution (see Figure 2-B).

Of particular importance is the independence of the genetic load and the mutational scheme (provided that mutation rates are pairwise symmetric; [4, 37]). Consequently, the load of Eq. (12) holds irrespectively of an organism’s degree of pleiotropy [37]: for instance, in a purely non-pleiotropic scenario where mutations would always affect one trait (and hence its fitness component) at a time, the genetic load is exactly the same than for any arbitrary complex pleiotropic scenario.

### 2.2 Unifying invariances in complexity-driven genetic load

In the special case of a uniform distribution of trait-dependent fitness components (*a* = *b* = 1) the WLE stationary genetic load obtained in Eq. (12) is:

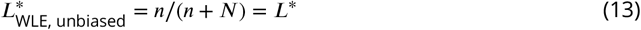

If we again assume a uniform distribution of fitness components and instead let organismal fitness be defined as *f*_*org*_(*r*) = 1 − *r*, we recover a version of Fisher’s geometric model that has previously appeared in the literature when *r* is given by the n-dimensional Euclidean distance *r*_FGM_ (**X**) between the phenotype **X** and the optimum [2, 4, 39]—albeit with a more explicit mutational scheme. Interestingly, the load for this version of Fisher’s geometric model is identical to the one we obtain here [2, 37]. This echoes previous results showing the invariance of the genetic load in modified versions of FGM with ellipsoidal rather than spherical fitness contours [12]. Together, these findings seem to suggest a broader unifying principle whereby the genetic load is invariant across a wide range of fitness landscape geometries.

#### 2.2.1 A sufficient condition for genetic load invariance

To elucidate why seemingly different fitness models yield the same evolutionary outcome, we can return to Eq. (10), in which the only factor that may differ from one fitness model to another is the density of genotypic fitness *ρ*(*f* ). Consequently, the genetic load obtained in Eq. (13), and indeed the entire stationary fitness distribution *ρ*^∗^(*f* ), remains invariant for all scenarios in which:

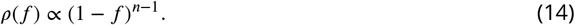

To see how this scaling arises in both WLE and FGM, it is helpful to work with fitness defects rather than fitness components (Figure 3A). To that end, we define the fitness defect associated with trait *d* as *X*_*d*_ = 1 −*F*_*d*_ . Since fitness components cannot exceed their optimum *F*_*d*_ = 1, each defect satisfies 0 ≤ *X*_*d*_ ≤ 1, and the optimal phenotype corresponds to (0, …, 0) in the space of defects. We then combine the trait-dependent fitness defects **X** = (*X*_1_, …, *X*_*n*_) into the organismal fitness defect *r* through a rule *r*_*org*_(**X**), which in turn relates to fitness as *f*_*org*_ (**X**) = 1 − *r*_*org*_(**X**). Accordingly, the density of fitness defects *ρ*(*r*) is simply related to the density of fitnesses *ρ*(*f* ) by *ρ*(*r*) = *ρ*(1 − *f* ). Through this lens, WLE can be naturally reinterpreted: fitness is fully dictated by the distance between the optimum and the most deleterious fitness component, so that *r*_WLE_ (**X**) = max(**X**), recovering *f*_WLE_ (**X**) = 1 − *r*_WLE_ (**X**) = 1 − max(**X**) = min(**F**).

**Figure 3.**
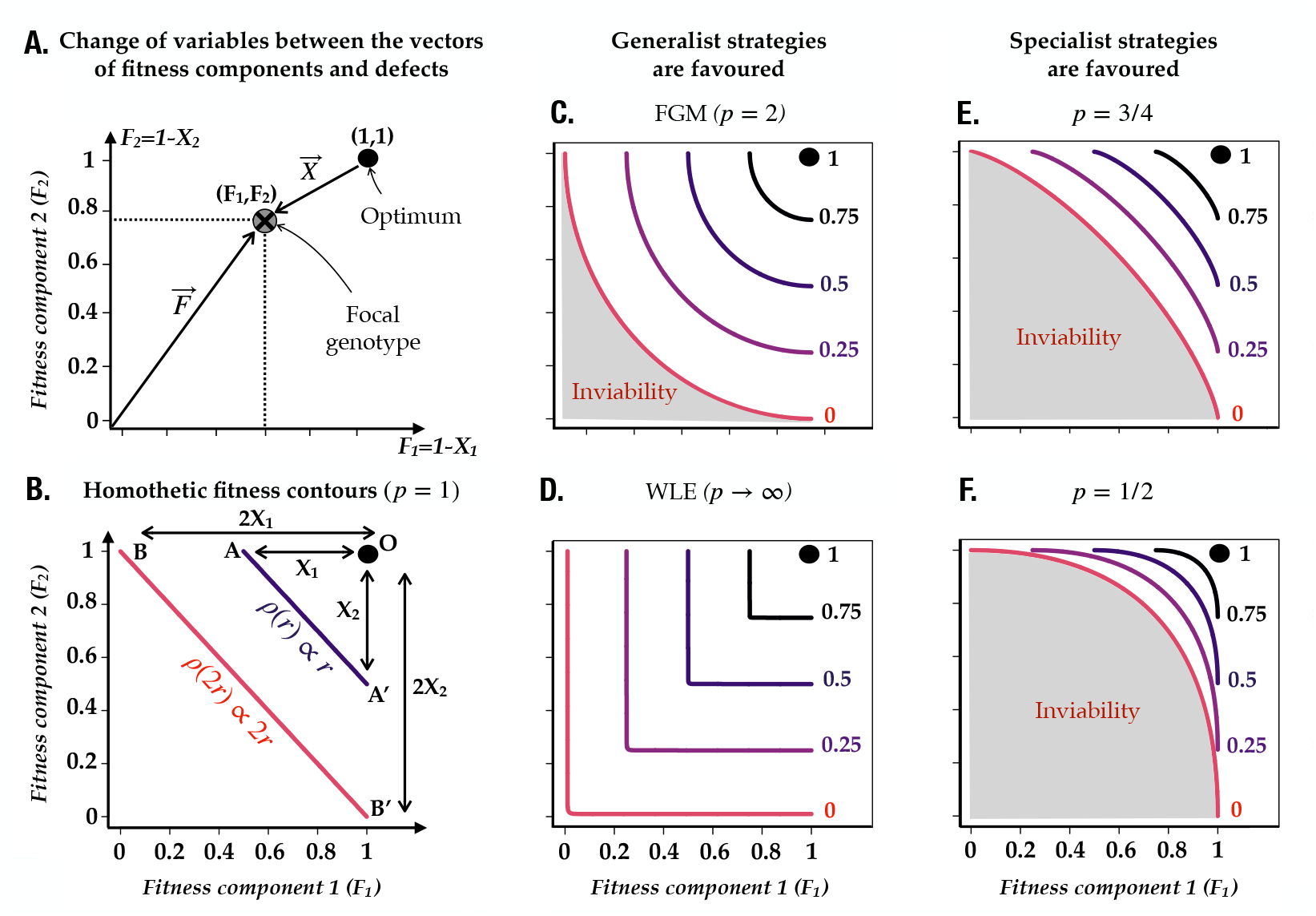
Fitness contours in a two-trait space. **A**. Change of variables used to interpret weakest link epistasis (WLE) in terms of the vector of fitness defects **X** = **1** − **F**. We restrict **F** to the unit cube (components ∈ [0, 1]), where the fitness optimum *f* = 1 is reached when all the *F*_*d*_ = 1 (here, *F*_1_ = *F*_2_ = 1). **B**. Illustration of the homothetic transformation principle behind the genetic load invariance when the organismal fitness defect is defined as *r*_*org*_ (**X**) = ||**X**||_1_ = *X*_1_ + *X*_2_ and fitness defects are uniformly distributed (*b* = 1). A homothety of ratio *k* is a transformation that maps any point at distance *X*_*i*_ from a choice of center (here *F*_1_ = *F*_2_ = 1) to a new point at distance *kX*_*i*_ (eg., A is sent to B here), preserving shapes and scaling all distances proportionally. Under uniform *ρ*(**X** (*b* = 1), *ρ*(*r*) is proportional to the *n* − 1-dimensional surface area of the equal fitness contours and scales as *r*^*n*−1^, where *n* = 2 traits in this example. **C–F**. Fitness contours (from *f* = 0 to *f* = 1, red-black gradient) resulting from fitness defined as *f* = 1 − *r*_*p*_(**X**), with *r*_*p*_(**X**) = ||**X**||_*p*_ denoting the genotypic fitness defect. Fitness defects are uniformly distributed (*b* = 1). The grey-shaded area represents combinations of fitness defects that are inviable, and expands as *p* decreases. **C–D**. Two ‘generalist’ strategies are shown, where improving both traits confers a higher fitness than improving a single one. The limit *p* → ∞ (panel D) recovers WLE (see Figure 1 for more details), while contours with *p* = 2 retains the weakest-link constraint, albeit in a softened form (panel C, canonical FGM with euclidean distance). **E–F**. Two ‘specialist’ strategies are shown (panel E: *p* = 2/3; Panel F: *p* = 1/2) where concentrating improvement on one trait is most efficient for increasing fitness.

Our goal is now to identify a broad class of rules *r*_*org*_(**X**), which, when combined with *f*_*org*_ = 1 − *r*_*org*_(**X**), result in *ρ*(*f* ) ∝ (1 − *f* )^*n*−1^. Working in fitness defects, the corresponding condition is:

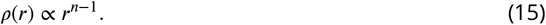

Figure 3 shows several rules *r*_org_ that satisfy this scaling (when *b* = 1). These include the additive rule 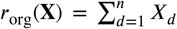 shown in panel B with diagonal equal-fitness contours, as well as Fisher’s geometric model *r*_FGM_ and weakest-link epistasis *r*_WLE_, shown in panels C and D, respectively, with circular and “L”-shaped equal-fitness contours. What all of these rules have in common is that they produce contours (i.e. level sets) that are rescaled versions of each other, expanding uniformly away from the optimum in what is known as a homothetic transformation (see Figure 3-B). Intuitively, this explains the *n* − 1 power scaling of the density, since *ρ*(*f* ) is proportional to the (*n* − 1 dimensional) surface area of the corresponding contour and, in *n*-dimensions, surface area scales as the (*n* − 1)-th–power of length.

Formally, a sufficient condition for this (*n* − 1) power scaling (when *b* = 1) is that *r*_*org*_ be a homogeneous function of degree one:

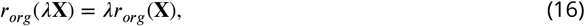

with *λ* > 0. This condition makes explicit the uniform scaling: for example, halving each trait-dependent fitness defect *X*_*d*_ also halves the organismal fitness defect *r*_*org*_, moving the system towards the optimum identically regardless of the background configuration **X**. More generally, this implies that the way trait-dependent fitness defects combine is independent of their overall scale (the effect of the rescaling factor *λ* is the same throughout the space of **X**). This has specific implications for mutational effects, since *r*_*org*_(**X**^′^)/*r*_*org*_(**X**) = *r*_*org*_(*λ***X**^′^)/*r*_*org*_(*λ***X**). If a mutation introduced in some background **X** reduces *r*_*org*_(**X**) to a fraction (e.g.1/3) of its current value, then the same mutation introduced in a background *λ***X** also reduces *r*_*org*_(*λ***X**) to the same fraction of its previous value.

Thus far, we have been working in the special case *b* = 1, so that genotypes are assumed to have trait-dependent fitness defects that are jointly uniformly distributed, i.e. *ρ*(**X**) ∝ 1. We next extend these results to a broader class of distributions of fitness defects. Notably, points lying on the same fitness contour may carry different weights determined by *ρ*(**X**), meaning there is no reason for the density *ρ*(*r*) to be strictly proportional to the surface area of the corresponding contour. In the Appendix B.1.2 we show that this differential weighting of points in the phenotypic space arises under a wide range of joint probability distributions *ρ*(**X**). Specifically, suppose that:

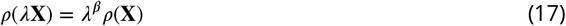

for all *λ* > 0 and some fixed value *β*. Technically, this says that we require that the density of fitness defects *ρ*(**X**) be a homogeneous function of degree *β*. More biologically, what this condition says is that the supply of lower fitness defect states changes in a consistent manner as the optimum is approached, so that, for instance, the density at **X**/2 is 2^*β*^ times smaller. Similar to the case for the organismal fitness defect, this means that all of the equal-density contours of the phenotypic distribution *ρ*(**X**) are scaled versions of each other, with yet more flexibility in the shape of the generating contour arising from the exponent *β*.

What we show in the Appendix B.1.2 is that for any *r*_*org*_ that is homogeneous of order 1 and any trait specific defect distribution *ρ*(**X**) that is homogeneous of degree *β*, then we have:

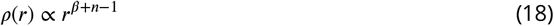

and hence, at stationarity, fitness is Beta(*N, β* + *n*)-distributed and results in a load given by (see section 2.1.2 for more details on the calculation):

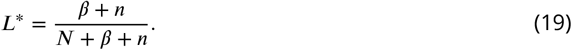

For instance, when the trait-dependent fitness components are independently Beta(1, *b*)-distributed, *ρ*(**X**) is a homogeneous function of degree *β* = *n*(*b* − 1) and we recover our previous expressions for the stationary fitness distribution and load (see supplemental appendix B.1.2).

This result shows that for fixed population size and a fixed degree of homogeneity for *ρ*(**X**), if *r*_*org*_ is homogeneous of degree 1, then the whole stationary distribution is invariant to the specific choice of how trait-dependent fitness defects combine and the structure of the neutral joint distribution of trait-dependent defects.

#### 2.2.2 Genetic load across a continuum of adaptation strategies

To more concretely explore the implications of the invariance in genetic load discussed in the previous section, we consider a family of choices for *r*_*org*_ that correspond to the *ℓ*^*p*^-norm of the vector of fitness defects:

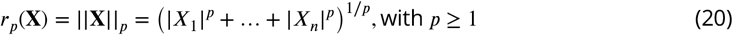

The *ℓ*^*p*^ norm provides a natural formalism for this purpose, since it satisfies the condition of homogeneity of degree 1 and recovers the Euclidean version of FGM (*p* = 2), additive trait specific fitness defects (*p* = 1), and also WLE version as a limit, where fitness is fully determined by the maximum fitness defect as *p* → ∞ (see Figure 3B-D)

Indeed there is an important ecological meaning behind different values of the parameter *p*. First, increasing *p* from the Euclidean version of FGM (*p* = 2) makes the model more and more dependent on the weakest link, which implies that intermediate values reflect a reality in between FGM and WLE: any trait improvement still enhances adaptiveness, albeit with diminishing returns, as often observed in biological systems [24]. In other words, higher *p* values reduce the ability to buffer maladaptive traits by limiting the effect (but not the supply) of compensatory mutations. Similarly, we can consider the impact of decreasing *p* from FGM (Figure 3E,F). Note that, in the cases where *p* < 1, ||*X*||_*p*_ is only a quasi-norm, since it fails to satisfy the triangle inequality, but it is still homogeneous of degree 1 and thereby satisfies the condition of Eq. (17). The case where *p* = 1 (also known as the Manhattan distance) represents the “symmetrical” situation, where fitness components/defects can buffer each other equally (e.g., adding 0.1 to to *X*_1_ and *X*_2_ is identical to adding 0.2 to *X*_1_, or to *X*_*n*_). When *p* falls under 1, the fitness contours tend to adopt more star-like shapes that favours putting all eggs in a single basket. In this “strongest link epistasis” (SLE) scenario, the fitness reward is higher for organisms that focus on improving one specific trait: because of the diamond shape of contours, the phenotypic distance to the optimum is much lower close to any given trait axis. Under this perspective, SLE can be seen as a model of specialisation, where all but specialist phenotypes would be driven out by natural selection.

A subtle but important implication of this model is that smaller values of p increasingly enlarge the region of trait-dependent fitness component space that results in inviable organisms (*r*_*p*_(**X**) > 1, implying negative fitness). For instance, in a two-trait example with *X*_1_ = 0.9 and *X*_2_ = 0.5, the genotype is inviable under the *r*_1_(**X**) and *r*_2_(**X**) rules (because both *r*_1_ > 1 and *r*_2_ > 1), yet retains viability fitness *f*_*org*_(**X**) = 0.1 under WLE. In our framework, these inviable combinations are still described by *ρ*(**X**) but make no contribution to the stationary distribution, which we consider to have support only on the biologically relevant interval from 0 to 1.

Turning to the issue of genetic load, as in Eq. (8), we assume again the fitness components to be independently Beta-distributed as *F*_*d*_ ∼ Beta(1, *b*). In this case, regardless of the parameter *p*, the *ℓ*^*p*^ norms are all homogeneous of degree 1, and hence the distribution of fitnesses at stationarity and corresponding genetic load *L*^∗^ are identical to that for WLE (Eqs. 9-12) (see APPENDIX B.1.2, as well as APPENDIX B.2.1 and B.2.2 for a more direct proof). The formula for the genetic load can even be further generalised by considering that fitness defects are not identically distributed and obey instead separate Beta(*b*_*d*_, 1) for *d* = 1 to *n*. Denoting mean trait-dependent fitness defect as 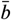 and assuming again 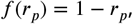, this yields (see APPENDIX B.1.2 and APPENDIX B.2.2):

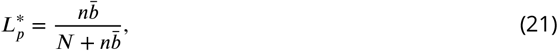

where 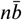 quantifies the effective maladaptive bias acting on fitness. This result also extends to fitness models with hierarchical genetic architecture, where fitness is determined by the *ℓ*^*p*^-norm of higher-level traits (e.g., metabolic pathways) and each of these traits is itself given by the 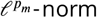 of underlying, lower level trait-dependent fitness defects, with *p*_*m*_ possibly differing across traits (see APPENDIX B.2.3 for the extension to modular inter-dependencies of traits).

## Discussion

Diminishing-returns effects are widespread in biology [19, 24, 26, 28]. They arise when improving a phenotypic trait (such as enzyme activity) no longer translates into increased performance of a function (such as metabolic flux) once another component becomes limiting. In some biological systems, phenotypes depend on the efficiency of every underlying component, as in multi-step biosynthetic pathways where failure of a single enzyme can fully prevent the formation of the final product (e.g., synthesis of anthocyanin pigments in plants). From a genetics perspective, such dependencies generate weakest-link epistasis (WLE), which can substantially distort the distribution of fitness effects.

Motivated by these considerations, we first determined the selection coefficient of mutations under WLE. Using this selection coefficient, we show how, in this fitness model, trait dimensionality greatly decreases the supply of advantageous phenotypes, which, in turn, limits the adaptive potential of organisms. Quantitatively, when fitness is determined by the weakest underlying trait, we establish that the the mean stationary genetic load carried by an organism scales as 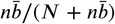, where 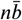 represents a compound maladaptive bias. A key feature of this expression is the symmetry between *n* and 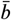: phenotypic complexity and intrinsically biased traits (towards low-fitness values) contribute symmetrically to the reduction of fitness.

Intriguingly, the genetic load under WLE can be shown to equal that of Fisher’s geometric model (FGM) for unbiased traits (*b* = 1), when fitness decays linearly with the Euclidean distance between the phenotype and the optimum [2, 37]. To elucidate why the complexity scaling of the load is preserved across diverse fitness landscapes despite the seemingly unfavourable genetic architecture imposed by WLE, we show that this arises due to the fact that the equal fitness contours in both cases consist of scaled copies of the same basic shape. We then generalise this scaling to allow for non-uniform joint distributions of trait-dependent fitness components.

To understand the effect of maladaptive biases across more interpretable forms of epistasis, we study the case where fitness defects to combine through arbitrary *ℓ*^*p*^ norms. This generalisation spans a continuum of architectures, including WLE (*p* → ∞), classical FGM (*p* = 2). This also naturally extends to combinations of *ℓ*^*p*^-norms that can capture biological modularity and hierarchical trait structure (eg., enzymes forming pathways that themselves combine to determine cell efficiency). Within this broader framework, the genetic load retains the *nb* scaling, demonstrating that neither the architecture of traits (e.g., modularity) nor the nature of epistasis alters the evolutionary stationary state: what matters is the number of independent phenotypic traits and their average bias toward low-fitness states.

In this work, we have defined fitness as *f*_*org*_(**X**) = 1 − *r*_*org*_(**X**). Earlier work has shown that the invariance of the genetic load can be further generalised by decoupling the rule that specifies the phenotypic ‘distance to the optimum’ *r*_*org*_(**X**) from the function *f*_*org*_(*r*) that maps this distance onto fitness, as is classically done in FGM [12]. For instance, a broad class of definitions for the phenotypic distance to the peak yields the same load 1 − ((*N* − 1)/*N*) ^*n*/*Q*^ when fitness is given by *f*_*org*_(*r*) = exp(−*r*^*Q*^).

Building on this invariance principle, these earlier approaches attempted top-down inferences of phenotypic dimensionality using the stationary genetic load [12, 21]. However, it remains to be determined whether biologically grounded fitness landscapes satisfy this homogeneity condition (even approximately), for instance in systems where fitness components combine multiplicatively across traits, a situation commonly encountered in the study of life-history traits [40, 41]. No less importantly, even when this condition holds, we have shown throughout this study that the symmetry between phenotypic dimensionality and maladaptive biases obscures the effects of complexity. As a consequence, stationary fitness alone cannot reliably disentangle these two sources of maladaptation unless maladaptive biases would be negligible.

This raises the question of whether there exists reliable empirical estimates of the magnitude of this (maladaptive) bias parameter *b*. Although the overwhelming predominance of non-functional proteins in random sequences has long been known [42], a more quantitative characterisation of biological quantities related to fitness components (eg., protein binding affinities) has only recently begun to emerge. These empirical distributions appear consistent with *b* values spanning several orders of magnitude [43, 44]. Crucially, this reinforces the idea that organisms may carry significant genetic loads, and strengthens the paradox of complexity: fitness gains associated with additional functions should come at the expense of increased maladaptation, that is, a larger difference between mean fitness and the optimal phenotype’s fitness for a given number of traits.

This perspective may also help clarify otherwise perplexing empirical findings. For instance, the vast majority of enzymes are only moderately efficient [45, 46], often to the point that their catalytic rates can be improved by several orders of magnitude [47]. Interpreting such patterns will require accounting in greater detail for the influence of genotype-phenotype maps on the distribution of fitnesses that would be realised under neutrality. More broadly, we believe that this framework paves the way for a more quantitative understanding of the outcome of evolution, such as the distribution of fitness effects of mutations. Likewise, it should help predict how biological systems evolve in the face of perturbations, for instance when environmental changes shift the location of the fitness peak, and how this depends on the underlying phenotype-fitness map.

## Acknowledgment

We thank Etienne Rajon, Djordje Bajic, Jason Wolf and Vitor Marquioni Monteiro for fruitful discussions that helped improve the exposition of the framework, and Dmitry Biba for the analogy with Liebig’s barrel. DMM acknowledges funding support from NIH grant R35GM133613 and additional funding from the Simons Center for Quantitative Biology at Cold Spring Harbor Laboratory.

## Supplementary Materials

### Supporting text A: Notation

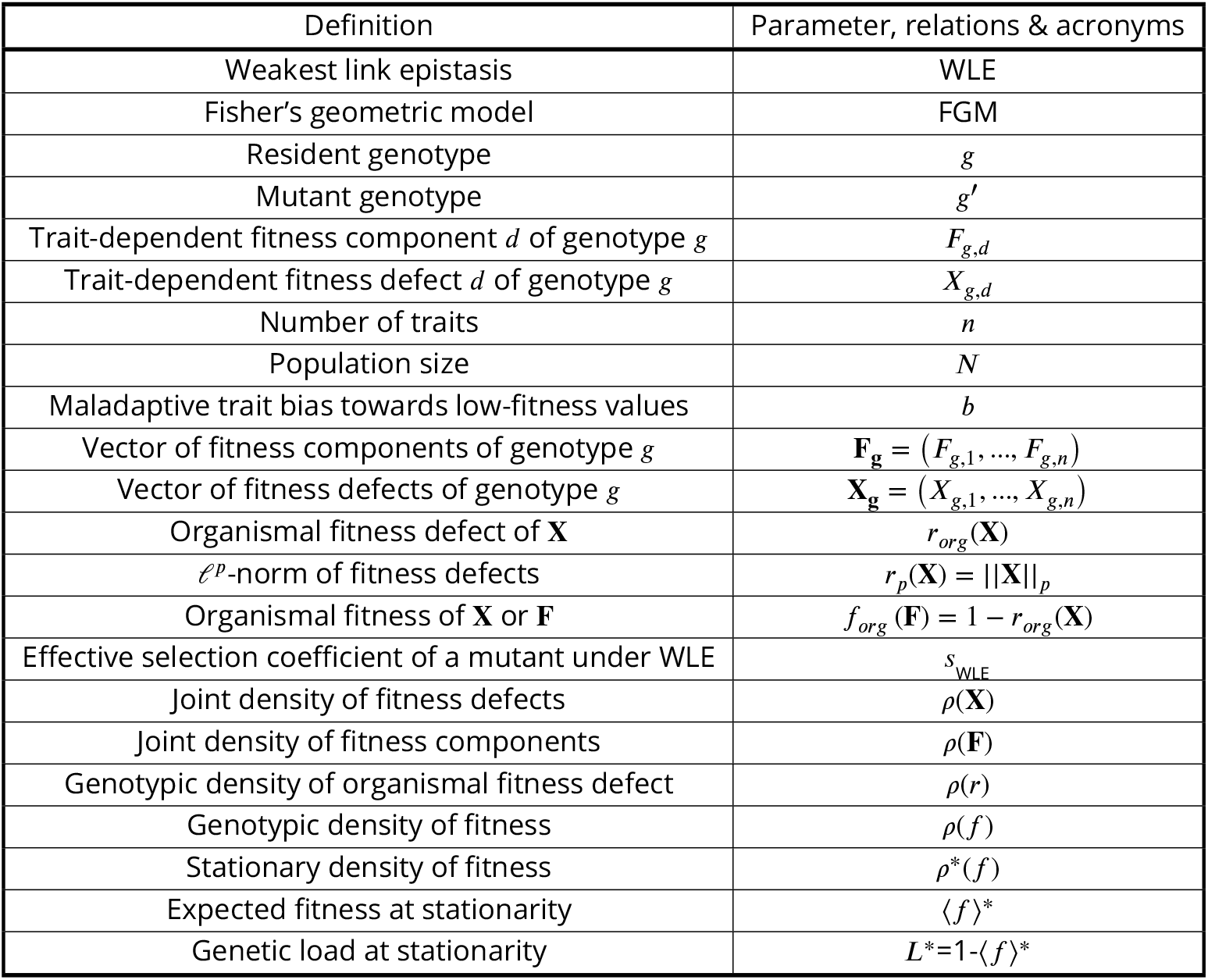

### Supporting text B: Mathematical appendix

#### B.1 Genetic load under various fitness models

##### B.1.1 WLE genetic load for the Kumaraswamy distribution

In this section, we relax the assumption that trait-dependent fitness components *F*_*d*_ are Beta(1, *b*) distributed. The first order statistics of a set of beta-distributed variables does not generally lead to a closed-form solution (when *a* ≠ 1), which precludes further analysis. The Kumaraswamy distribution 𝒦*umar*(*a, b*), (with shape parameters *a* and *b*) is a Beta-like distribution with compact support ([35]; see Figure 4 for the shape of these distributions) for which the first-order statistic has a closed-form regardless of parameters *a* and *b*: if the vector of fitness components **F** contains *n* random variables *F*_*d*_ ∼ 𝒦*umar*(*a, b*), then the distribution of their minimum is also Kumaraswamy-distributed [35, 36], such that :

**Figure 4.**
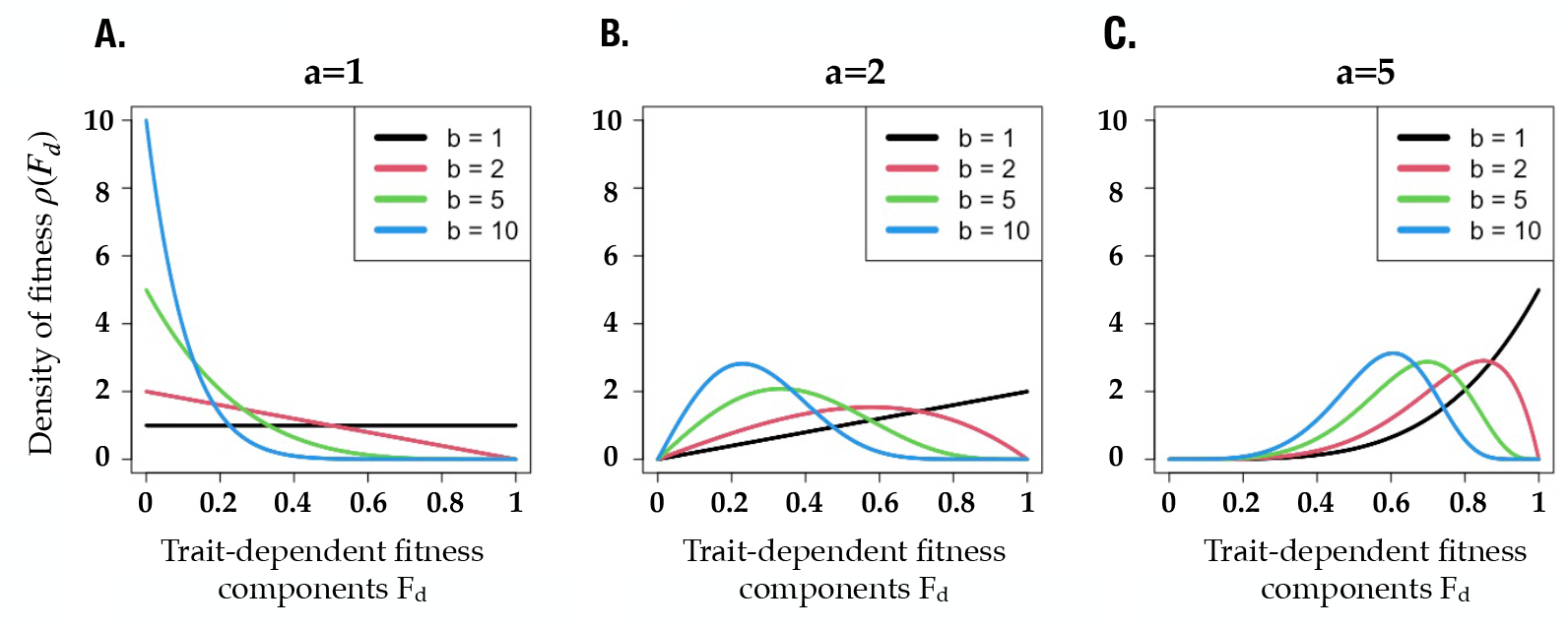
Shape of the genotypic distribution of trait-dependent fitness components when they are identically Kumaraswamy-distributed, *F*_*d*_ ∼ ℬ*umar*(*a, b*). Fitness components are bounded between 0 and 1 with their contribution to fitness being maximal at 1. Overall, the parameters *a* and *b* modify the genotypic distribution of trait values (i.e., prior to selection), thereby creating a bias toward deleterious traits (when *b* > *a*) or toward advantageous ones (when *a* > *b*). **A**. In the case where *a* = 1, *F*_*d*_ follows a Beta-distribution *F*_*d*_ ∼ ℬ*eta*(1, *b*) as modeled in the main document. The special case *a* = *b* = 1 coincides with a uniform distribution, and thus with the absence of bias at the trait level, whereas *b* = 2 leads to a linear decrease of the density of fitness component values (higher weight on low-fitness trait values). For larger *b*, the density is highly biased towards deleterious values. **B**–**C**. Increasing the parameter *a* relaxes the bias towards deleterious traits (*a* = 2, panel B), to the point where high fitness traits become over-represented (especially when *a* = 5, panel C).

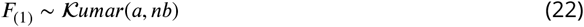

This arises because the influence of the parameter *a* is slightly modified compared to the Beta-distribution, as reflected by its probability density function. Specifically, the distribution of *F*_*d*_ when *F*_*d*_ ∼ 𝒦*umar*(*a, b*) is given by:

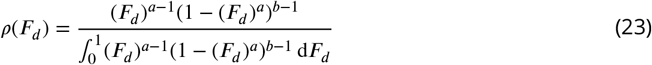

where the difference with the Beta-distribution is in the term (1 − (*F*_*d*_ )^*a*^), which would simply be (1 − *F*_*d*_ ) in a Beta-distribution.

Under weakest link epistasis, fitness is determined by the minimum of fitness components. Hence, Eq. (22) corresponds to the genotypic distribution of fitness *ρ*_WLE_ (*f* ) in the absence of selection.

Assuming a haploid population and a Moran evolutionary process, we can use Eq. (10) to derive the distribution of fitness at evolutionary stationarity:

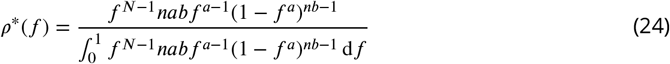

Fitness at evolutionary stationarity is thus given by a ratio of raw moments (see main text for explanations) that are analytically tractable for the Kumaraswamy distribution [36]:

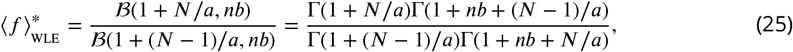

where ℬ (*y*) and Γ(*y*) denote the Beta and Gamma functions. From Stirling’s formula, we have the approximation Γ(*y*+*s*) ≃ *y*^*s*^Γ(*y*) when *y* is large enough, with which we can rewrite fitness at stationary state as:

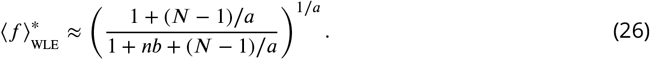

We can rewrite Eq. (26) as (1 − *y*)(1/*a*), where *y* = *nb*/ (1 + *nb* + (*N* − 1)/*a*)) is small when *nba* ≪ *N*. Using Taylor expansions, we can then simplify Eq. (25) so that in the haploid Moran scenario, we have:

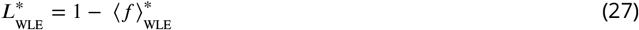

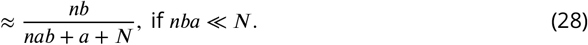

As a first order approximation, we could neglect all terms but *N* in the denominator, but keeping the *nba*+*a* term both improves the approximation and helps gain a better intuition of the role played by *n,a* and *b* when their product gets close to *N*.

From this expression, we first recover that population size *N* reduces the genetic load. This expression then shows that dimensionality *n* and the parameter *b* tend to increase the genetic load in a purely exchangeable way. In the main document, we use the power version of the Beta distribution where *a* = 1, such that *b* directly represents an intrinsic bias toward low-fitness traits. With the Kumaraswamy distribution (or a Beta-distribution where *a* ≠ 1, this bias is better captured by the ratio *b*/*a*. Accordingly, while *b* keeps its influence on inflating low fitness genotypes, the parameter *a* acts as a counter-bias. Hence, it reduces the genetic load by providing evolution with a large(r) supply of high fitness genotypes. Notably, this result is robust to a broad range of fitness-component distributions, since the Kumaraswamy distribution can approximate diverse shapes, including Gaussian- and exponential-like distributions (see Figure 4). Overall, *a* has a compound influence that is partly akin population size, partly limiting the influence of the maladaptive bias *nb*.

Logically, this formula for the genetic load becomes exact when *a* = 1, as Γ(*y*+1) = *y*Γ(*y*). Starting back from Eq. (26), we can thus write the genetic load exactly as the expression of the main document, because the Kumaraswamy distribution ℬ*umar*(1, *b*) reduces to a Beta-distribution *Beta*(1, *b*) in this case (which also explains why the Beta distribution with parameter *a* = 1 has a tractable first order statistics, as we claim in the main document):

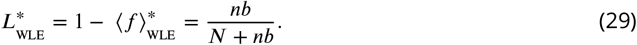

##### B.1.2 Invariance of stationary fitness distribution

Here we derive the stationary fitness distribution when both the organismal fitness and the genotypic density are homogeneous functions of the trait specific fitness defects. First, we prove the following technical lemma.

###### Lemma 1.

*Let g be a homogeneous function of degree α* > 0 *and let h be a homogeneous function of degree β, both defined on the positive orthant of* ℝ^*n*^. *Let*

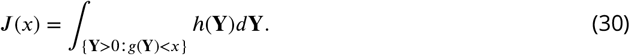

*Then J* (*x*) *is a homogeneous function of degree*

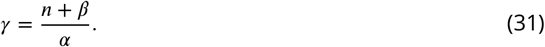

*Proof*. Consider

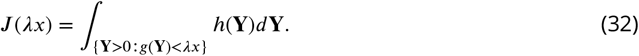

We will evaluate this integral by using the change of variables **Y** = *λ*^1/*α*^**Z**. This change of variables is an invertible linear transformation defined by the matrix *λ*^1/*α*^I where I is the identity matrix and hence the determinant of its Jacobian is simply |*λ*^1/*α*^I| = *λ*^*n*/*α*^ for all **Z**. Turning to the region of integration, *g*(**Y**) < *λx* if and only if *g*(*λ*^1/*α*^**Z**) < *λx*, but *g* is homogeneous of degree *α* and so *g*(*λ*^1/*α*^**Z**) = *λ g*(**Z**). Thus, *g*(**Y**) < *λx* if and only if *g*(**Z**) < *x*. Likewise, since **Y** is a positive multiple *λ*^1/*α*^**Z** of **Z, Y** > 0 if and only if **Z** > 0.

Putting this all together, we have

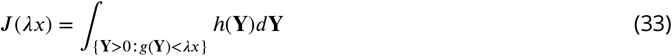

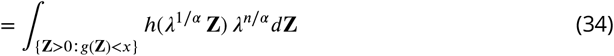

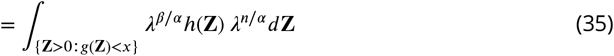

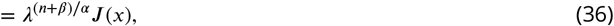

where in line 35 we used the fact that *h* is homogeneous of degree *β*. Thus *J* (*λx*) = *λ*^(*n*+*β*)/*α*^*J* (*x*) meaning that *J* is homogeneous of degree (*n* + *β*)/*α* as required.

We can now characterise the genotypic density of organismal fitness defect *ρ*(*r*) induced by a homogeneous function *r*_*org*_(**X**) mapping trait-dependent fitness defects to organismal fitness defects when the genotypic density of trait-dependent fitness defects is itself also a homogeneous function.

###### Proposition 1.

*Suppose the organismal fitness defect r*_*org*_(**X**) *is homogeneous of degree 1, and ρ*(**X**) *is homogeneous of degree β. Then ρ*(*r*) ∝ *r*^*n*+*β*−1^.

*Proof*. Let

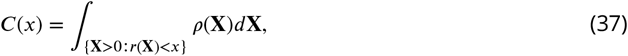

so that *ρ*(*r*) is proportional to *C*^′^(*x*). By Lemma 1, *C*(*x*) is a homogeneous function of degree *n* + *β*, and hence its derivative *C*^′^(*x*) is a homogeneous function of degree *n* + *β* − 1, as required.

The stationary distribution of fitness itself, and hence the expected load at stationarity, then follow as an immediate consequence of Equation 10.

###### Corollary 1.

*Suppose the organismal fitness defect function r*_*org*_(**X**) *is homogeneous of degree 1 and ρ*(**X**) *is homogeneous of degree β. Then for a Moran population of size N (as we used when deriving the WLE load) the stationary distribution of f is* ℬ*eta*(*N, β* + *n*) *distributed where n is the number of traits. Moreover, the load is given by:*

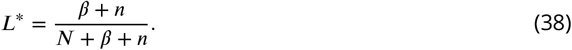

*Proof*. By Proposition 1, *ρ*(*r*) ∝ *r*^*n*+*β*−1^ and the density of *f* is simply a reflected version of the density of *r*, since *f* = 1 − *r*, so that we have *ρ*(*f* ) ∝ (1 − *f* )^*n*+*β*−1^. Plugging these values into Equation 10, we find that the stationary distribution of *f* has density

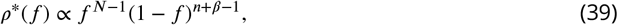

which we recognize as the density of a ℬ*eta*(*N, β* +*n*) distribution. Since the mean of a ℬ*eta*(*N, β* +*n*) distribution is *N*/(*N* + *β* + *n*), the stationary load *L*^∗^ is

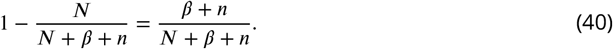

**Corollary 2**. *Suppose each single-trait defect X*_*d*_ *is homogeneous of degree b*_*d*_ − 1, *with b*_*d*_ *denoting the intrinsic maladaptive bias of trait d. Then for a Moran population of size N the stationary distribution of f is* 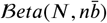 *distributed where n is the number of traits. Moreover, if* 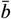 *denotes the mean maladaptive bias across traits, the load is given by:*

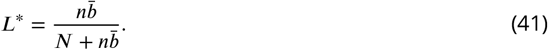

*Proof*. If the probability distribution of each trait *ρ*(*X*_*d*_ ) is homogeneous of degree *b*_*d*_ − 1, the joint distribution is given by the product of the marginal distributions. It is thus homogeneous of degree 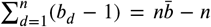. Replacing by 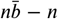 in the previous corollary completes the proof for this second corollary.

In the next section, we use the special case of the *ℓ*^*p*^ norm to relate this reasoning to a full probabilistic approach.

#### B.2 Beta-distributed variables and the *ℓ*^*p*^ norm

##### B.2.1 Distribution of the *ℓ*^*p*^ norm of Beta-distributed variables

###### Proposition 2.

*Consider a set of n independent variables X*_1_, *X*_2_, …, *X*_*d*_, …, *X*_*n*_ *each distributed as X*_*d*_ ∼Beta(*a*_*d*_, 1) *with a*_*d*_ > 0. *Conditional on* ||**X**||_*p*_ ≤ 1, *the distribution obeyed by the ℓ*^*p*^ *norm follows:*

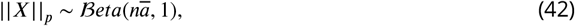

*with* ||*X*||_*p*_ = (|*X*_1_|^*p*^ + … + |*X*_*n*_|^*p*^)^1/*p*^ *and* 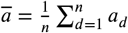.

This result is well established for *X*_*d*_ that are uniformly distributed [48]. Below, we extend it to the case of Beta-distributed variables.

###### Lemma 2.

*To prove the proposition in Eq. (*42*), we first need to show the following lemma. If X*_*d*_ ∼ ℬ*eta*(*a*_*d*_, 1) *with a*_*d*_ > 0, *then* :

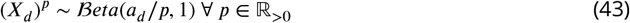

*Proof*. To do so, we need to determine the probability density function 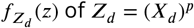. Using a simple change of variable and the chain rule (for derivatives), it is possible to write the probability density function 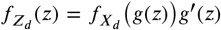 provided *x* = *g*(*z*) is a strictly increasing differentiable function. Applying this to the function with *X*_*d*_ ∼ ℬ*eta*(*a*_*d*_, 1), we have 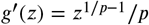 This implies:

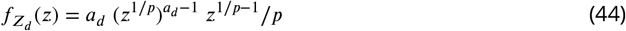

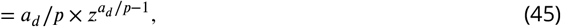

which completes the proof since this expression is exactly the probability density function of a Beta distribution ℬ*eta*(*a*_*d*_ /*p*, 1). From our premises, it is besides possible to write *q* = 1/*p*, which immediately yields the useful corollary according to which, if *X*_*d*_ ∼ ℬ*eta*(*a*_*d*_ /*q*, 1) with *a*_*d*_ > 0:

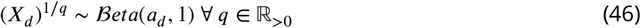

The following lemma can be derived from the aggregation property of the Dirichlet distribution by aggregating the first *n* variables of Dir(*α*_1_, …, *α*_*n*_, 1), however here for completeness we provide a direct proof.

###### Lemma 3.

*Consider a set of n variables Y*_*d*_ ∼ ℬ*eta*(*α*_*d*_, 1) *with α*_*d*_ > 0. *Conditioning on* 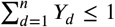, *we can show that:*

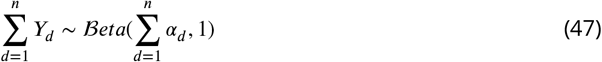

*Proof*. We start with two independent variables *Y*_1_ ∼ ℬ*eta*(*α*_1_, 1) and *Y*_2_ ∼ ℬ*eta*(*α*_2_, 1) that have compact support on [0, 1], with *α*_1_, *α*_2_ > 0. The probability density function for the sum *S* = *Y*_1_ + *Y*_2_ is given by the convolution:

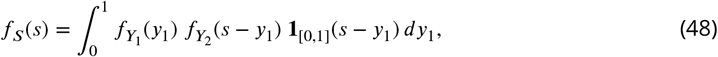

where **1**_[0,1]_ denotes the indicator function that enforces variables to live on their support. Because both variables *Y*_1_ and *Y*_2_ take values between 0 and 1, their density vanishes outside this interval. Consequently, *y*_2_ = *s* − *y*_1_ ∈ [0, 1] has to be positive. This is enforced by the term **1**_[0,1]_(*s* − *y*_1_), and implies that the integrand is supported only for *y*_1_ < *s*; otherwise, this would require *y*_2_ < 0. We can thus write:

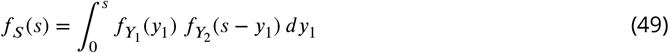

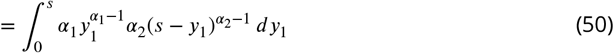

Exchanging *y*_1_ for *us*, we have *dy*_1_ = *sdu* and we can rewrite Eq. (50) as:

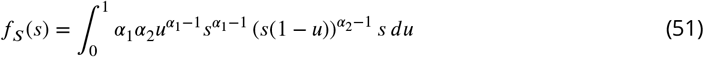

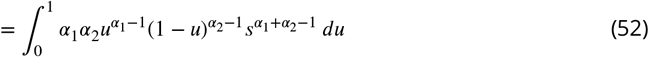

We can now include the truncation following which the density of the sum is *f*_*S*_(*s* > 1) = 0 when the sum of *Y*_1_ and *Y*_2_ is greater than 1. The probability density function must thus be renormalised over the simplex. This can be written as:

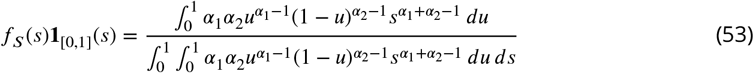

where normalisation occurs via the denominator. Since *u* is not a function of *s*, the integration over *u* is independent of *s*. Consequently, all terms that depend on *u* cancel out in the expression, which reduces to:

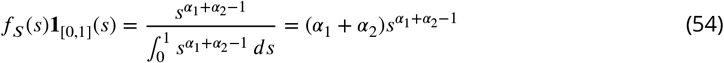

By definition of the Beta distribution, we have shown through Eq. (54) that the sum of *Y*_1_ and *Y*_2_ obeys *S* ∼ ℬ*eta*(*α*_1_ + *α*_2_, 1).

Remarking that the second parameter of the distribution is conserved by the summation, we can extend this property by combining the sum *S* with a third variable *Y*_3_, so that (*S*+*Y*_3_) ∼ ℬ*eta*((*α*_1_+ *α*_2_) + *α*_3_, 1). Extending it further to *n* variables demonstrates the above lemma.

We can now combine Lemmas (2) and (3) to demonstrate the proposition (2).

*Proof*. First, including Lemma (2) into Lemma (3) yields:

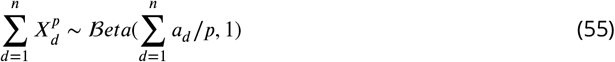

when we condition on 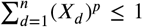. Second, we can notice that this latter requirement is equivalent to ||**X**||_*p*_ ≤ 1 because variables *X*_*d*_ ∈ [0, 1], and hence |*X*_*d*_ | = *X*_*d*_ . Therefore, using Lemma (2) again on the sum 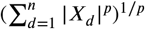 concludes the proof by canceling out the influence of *p* in the distribution of Eq. (55) and making it equal to Eq. (42).

#### B.2.2 *ℓ*^*p*^ norm and the distribution of fitness defects

We can now apply the property given by condition in Eq. (42) to the vector of fitness defects. We have defined this vector as **X**, where *X*_*d*_ ∼ Beta(*b*_*d*_, 1). From Eq. (42), we have:

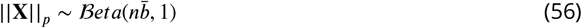

With *r*_*p*_ = ||**X**|| _*p*_ and *f* = 1 − *r*_*p*_, we then immediately recover that:

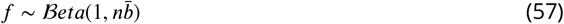

owing to reflection symmetry. This is equivalent to Eq. (9) except it now relies on the average maladaptive bias 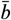 rather than *b*. As an immediate consequence, the genetic load takes the form of Eq. (12), with 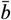 instead of *b*.

##### B.2.3 Distribution of the sum of Beta-distributed *ℓ*^*p*^-norms conditioning on ≤ 1, and connection to modularity

###### Distribution of the sum of *ℓ*^*p*^**-norms conditional on** ||**X**||_*p*_≤ 1

There is another useful consequence of the fact that the *ℓ*^*p*^ norm of a vector of independent Beta distributed variables *X*_*d*_ ∼ ℬ*eta*(*a*_*d*_, 1) remains Beta-distributed when we condition on ||**X**||_*p*_ ≤ 1: we can now apply this result to determine the distribution followed by the combination of *ℓ*^*p*^ norms themselves.

First, consider M subsets of *n*_*m*_ independent random variables following Beta-distributions:

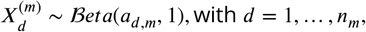

and the group-level measure of each subset as the 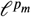 norm:

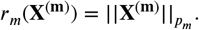

Conditional on 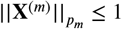, our prior results imply that:

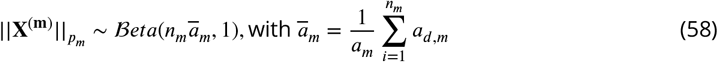

From that point, we can now determine the distribution followed by the sum of several *ℓ*^*p*^-norms, and by taking the *ℓ*^*p*^-norm of underlying *ℓ*^*p*^-norms.

###### Corollary 3.

*Consider M subsets of vectors* **X**^(**m**)^ *where m* = 1, …, *M, such that* :

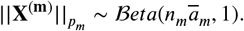

*Now, consider the set of these* 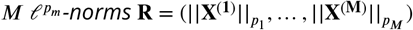 *of order p*_*m*_. *If the* 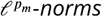 *are combined through a higher-level ℓ*^*p*^*-norm, the resulting distribution* ||**R**||_*p*_ *of is still Beta-distributed. Importantly, this property does not depend on the specific values of the exponents p*_*m*_, *or the number of hierarchical layers*.

*Proof*. Consider a vector containing *M* group-level norms:

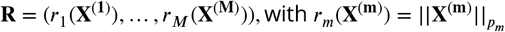

Conditioning on ||**R**||_*p*_ ≤ 1, we can apply property (2), and the *ℓ*^*p*^ norm is again Beta-distributed according to:

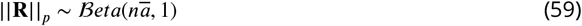

with:

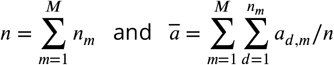

Consequently, the distribution of the combined *ℓ*^*p*^ norm is completely independent of the number of layers, the choice of the orders *p*_*m*_ at each layer, and the final order *p*.

###### Biological modularity and the genetic load under maladaptive biases

We can use this property to clarify how the genetic load invariance relates to the architecture of traits. An organism may consist of multiple functional modules, within which lower-level traits combine in different ways. For example, the size of an organism may arise as the sum of the sizes of smaller components (eg., bones in a body), which corresponds to the taxicab *ℓ*^1^ norm. conversely, a metabolic rate or the synthesis of a pigment may require all lower-level components to contribute jointly (eg., several glycolytic enzymes are involved to produce ATP), which corresponds to the maximum *ℓ*^∞^ norm in the space of defects. These higher-level traits may then combine to prescribe fitness. Eq. (59) demonstrates that neither modular nor hierarchical architectures of traits modify the genetic load. This latter simply depends on the total number of traits and on the average maladaptive bias of genotypes toward producing low-fitness trait values.

